# nNOS-expressing neurons in the ventral tegmental area and substantia nigra pars compacta

**DOI:** 10.1101/251843

**Authors:** Eleanor J Paul, Eliza Kalk, Kyoko Tossell, Elaine E. Irvine, Dominic J. Withers, James Leiper, Mark A Ungless

**Affiliations:** MRC London Institute of Medical Sciences (LMS), Du Cane Road, London W12 0NN, UK; Institute of Clinical Sciences (ICS), Faculty of Medicine, Imperial College London, Du Cane Road, London W12 0NN, UK

## Abstract

GABA neurons in the ventral tegmental area (VTA) and substantia nigra pars compact (SNc) play key roles in reward and aversion through their local inhibitory control of dopamine neuron activity and through long-range projections to several target regions including the nucleus accumbens. It is not clear if some of these GABA neurons are dedicated local interneurons or if they all collateralize and send projections externally as well as making local synaptic connections. Testing between these possibilities has been challenging in the absence of interneuron-specific molecular markers. We hypothesised that one potential candidate might be neuronal nitric oxide synthase (nNOS), a common interneuronal marker in other brain regions. To test this, we used a combination of immunolabelling (including antibodies for nNOS that we validated in tissue from nNOS-deficient mice) and cell-type-specific virus-based anterograde tracing in mice. We show that nNOS-expressing neurons in the parabrachial pigmented (PBP) part of the VTA and the SNc are GABAergic local interneurons, whereas nNOS-expressing neurons in the Rostral Linear Nucleus (RLi) are mostly glutamatergic and project to a number of regions, including the lateral hypothalamus, the ventral pallidum, and the median raphe nucleus. Taken together, these findings indicate that nNOS is expressed by neurochemically- and anatomically-distinct neuronal sub-groups in a sub-region-specific manner in the VTA and SNc.

## Introduction

Around one third of neurons in the ventral tegmental area (VTA) and substantia nigra pars compact (SNc) are γ-Aminobutyric acid (GABA)-ergic (Olson & Nestler, 2007; Nair-Roberts *et al.*, 2008). These neurons make local, inhibitory synaptic connections with dopamine neurons and their activation can drive conditioned place aversion and reduce food consumption (Omelchenko & Sesack, 2009; Tan *et al.*, 2012; van Zessen *et al.*, 2012). In addition, they send long-range axonal projections to several target regions, including the nucleus accumbens where they can regulate associative learning (Brown *et al.*, 2012; Taylor *et al.*, 2014). It is not clear if a subset of these GABA neurons are dedicated local interneurons or if they all collateralize and send projections externally as well as making local synaptic connections. Testing between these possibilities has been challenging in the absence of interneuron-specific molecular markers. Indeed, of the cardinal interneuron markers used to identify and selectively target sub-populations of interneurons in other regions of the brain, most are either not expressed in either the VTA or SNc, or are also expressed by sub-groups of dopamine neurons (e.g., somatostatin, cholecystokinin, vasoactive intestinal peptide, neuropeptide Y, parvalbumin, and calretinin (Hokfelt *et al.*, 1980; Seroogy *et al.*, 1988; Seroogy *et al.*, 1989; Rogers, 1992; Isaacs & Jacobowitz, 1994; Liang *et al.*, 1996; Gonzalez-Hernandez & Rodriguez, 2000; Klink *et al.*, 2001; Lein *et al.*, 2007; Olson & Nestler, 2007; Dougalis *et al.*, 2012; Merrill *et al.*, 2015). One potential candidate, however, is neuronal nitric oxide synthase (nNOS). nNOS is a member of the nitric oxide synthase family of enzymes that catalyse the synthesis of nitric oxide (NO) from L-arginine (Knowles *et al.*, 1989; Garthwaite, 1991). In the nervous system NO acts as a gaseous transmitter that can move rapidly across plasma membranes in anterograde and retrograde directions (Garthwaite & Boulton, 1995; Wang & Marsden, 1995). In several brain regions nNOS is selectively expressed by specific types of GABAergic interneurons (Klausberger & Somogyi, 2008; Tepper *et al.*, 2010a). Although several reports indicate that nNOS is expressed sparsely in the VTA and/or the SNc, there are discrepancies regarding the extent of its expression, which subregions it is expressed in, and the degree of colocalisation with tyrosine hydroxylase (the rate limiting enzyme in dopamine synthesis that is most commonly used to identify dopamine neurons) (Vincent & Kimura, 1992; Rodrigo *et al.*, 1994; Gonzalez-Hernandez & Rodriguez, 2000; Backes & Hemby, 2003; Klejbor *et al.*, 2004; Gotti *et al.*, 2005; Cavalcanti-Kwiatkoski *et al.*, 2010; Mitkovski *et al.*, 2012). We hypothesised that some of these discrepant findings may have arisen because of non-specific immunolabelling. To address this directly, we tested three different nNOS antibodies for reliable immunolabelling in the VTA and SNc, using tissue from nNOS-deficient mice as a control. This allowed us to establish that only one of these antibodies exhibited reliable immunolabelling in the VTA and SNc. Using this antibody, combined with cell-type-specific viral-based anterograde axonal tracing, we found that nNOS is expressed by several distinct sub-groups of neurons in the VTA and SNc, including GABAergic interneurons and glutamatergic projection neurons.

## Materials and Methods

### Animal maintenance and breeding

C57Bl/6NCrl (RRID: IMSR_CR:027; WT) mice were purchased from Charles River. nNOS-deficient (RRID: IMSR_JAX:002986), NOS1Cre (RRID: IMSR_JAX:017526), VGATCre (vesicular GABA transporter; RRID: IMSR_JAX:016962), and RiboTag (RRID: IMSR_JAX:011029) mice were purchased from the Jackson Laboratory. Mice heterozygous for VGATCre (VGATCre -/+) were crossed with mice homozygous for RPL22^HA^ (RiboTag +/+) producing VGATCre -/+ RiboTag -/+ offspring (VGATCre:RiboTag). NOS1Cre mice were heterozygous. All breeding and experimental procedures were conducted in accordance with the Animals (Scientific Procedures) Act of 1986 (UK). All mice were maintained in social groups of 2-4, where possible, with appropriate environmental enrichment (e.g., bedding and tunnels). They were kept in rooms at a constant temperature and maintained on a 12 h light/dark cycle. They were fed on standard rodent chow and water ad libitum.

### Tissue fixation and preparation

C57Bl/6NCrl, nNOS-deficient, VGATCre:RiboTag, or NOS1Cre mice were anaesthetised under isoflurane (4 %) and given a lethal intraperitoneal (IP) injection of pentobarbital (100mg/ml; Euthatal). They were transcardiallly perfused with 50 ml of ice cold phosphate buffered saline (PBS) followed by 50-100 ml of 4 % paraformaldehyde (PFA; Sigma Aldrich) in PBS. When fixed, the brains were removed and placed in 10 ml of 4 % PFA for 1 h post-fixation at room temperature. After 3 washes in PBS, brains were placed in 30 % sucrose (Sigma Aldrich) dissolved in PBS for cryo-protection, and kept at 4 °C for 24-48 h. Subsequently, all brains were embedded in optimal cutting temperature (OCT) medium and snap frozen in isopentane (2-methlybutane) at -55 °C. All tissue was then stored at -80 °C until sectioning.

### Immunocytochemistry

All immunolabelling was conducted on tissue from mice aged 8-12 weeks old. Brains were sectioned using a Leica CM1800 cryostat (Leica Microsystems, Germany). Coronal sections (30 µm) were taken from the midbrain, or from the whole brain in the case of Nos1Cre mice. Free floating sections were washed in PBS for 10 min at room temperature. Following this, they were blocked in 6 % normal donkey serum (NDS) in 0.2 % Triton-x in PBS (PBSTx) for 60 mins at room temperature. Primary antibodies (Table 1) were diluted in 2 % donkey serum in PBSTx and sections were incubated in the primary antibody solutions overnight at 4 °C. Sections were washed (3 x 10 mins) in PBS at room temperature. Secondary antibodies (Table 2) were diluted in 2 % donkey serum 0.2 % PBSTx. Sections were incubated in secondary antibody solution for a minimum of 1.5 h at room temperature. They were then washed (3 x 10 mins) in PBS. Stained sections were mounted onto glass microscope slides and when dry were cover-slipped using VectaShield mounting medium (Vector Laboratories). SNc and VTA regions were determined using tyrosine hydroxylase (TH) expression. Region outlines were traced from Franklin and Paxinos (2008).

**Table 1.**
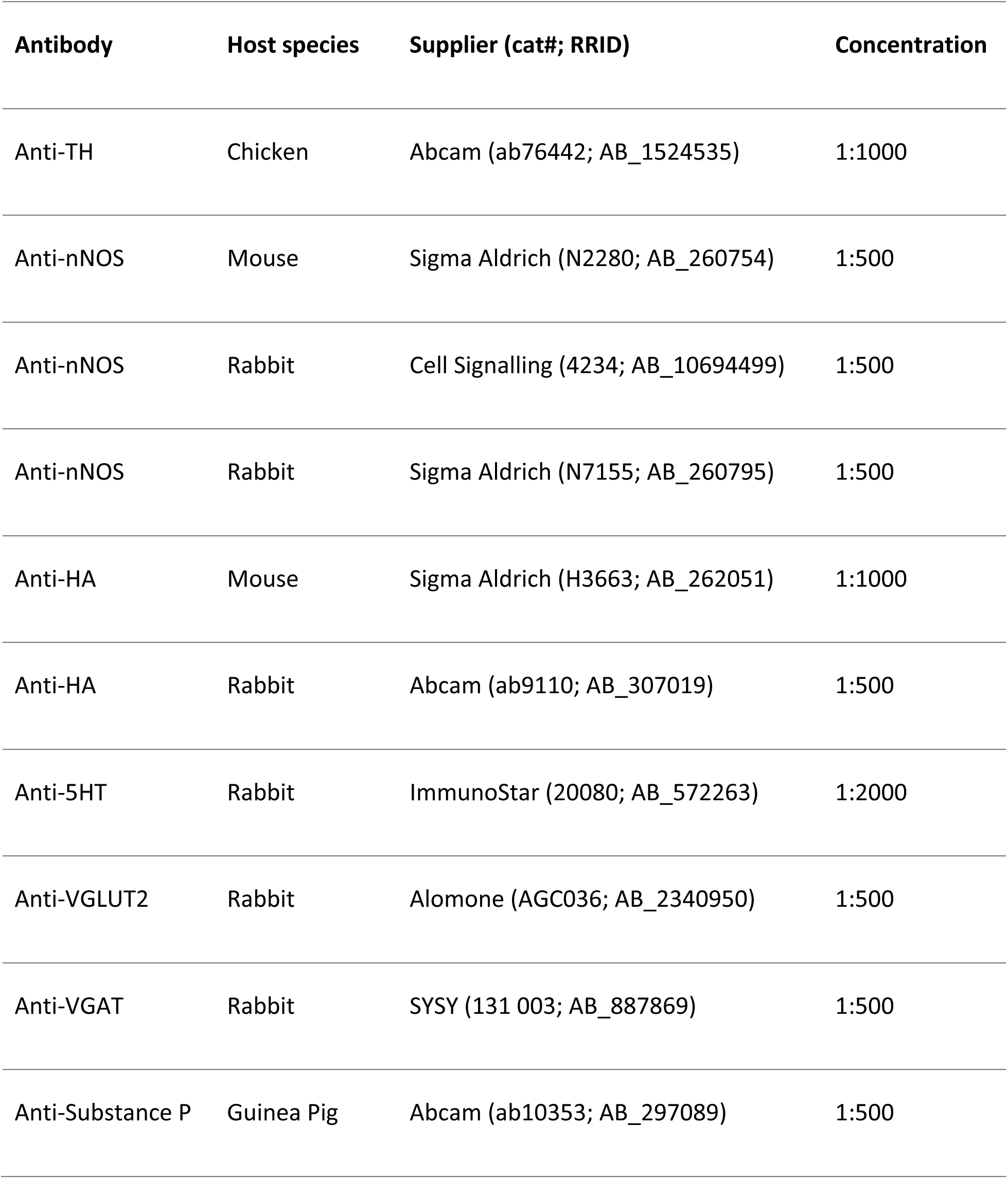
**Primary antibodies**

| **Antibody** | **Host species** | **Supplier (cat#; RRID)** | **Concentration** |
| --- | --- | --- | --- |
| Anti-TH | Chicken | Abcam (ab76442; AB_1524535) | 1:1000 |
| Anti-nNOS | Mouse | Sigma Aldrich (N2280; AB_260754) | 1:500 |
| Anti-nNOS | Rabbit | Cell Signalling (4234; AB_10694499) | 1:500 |
| Anti-nNOS | Rabbit | Sigma Aldrich (N7155; AB_260795) | 1:500 |
| Anti-HA | Mouse | Sigma Aldrich (H3663; AB_262051) | 1:1000 |
| Anti-HA | Rabbit | Abcam (ab9110; AB_307019) | 1:500 |
| Anti-5HT | Rabbit | ImmunoStar (20080; AB_572263) | 1:2000 |
| Anti-VGLUT2 | Rabbit | Alomone (AGC036; AB_2340950) | 1:500 |
| Anti-VGAT | Rabbit | SYSY (131 003; AB_887869) | 1:500 |
| Anti-Substance P | Guinea Pig | Abcam (ab10353; AB_297089) | 1:500 |

## Microscopy

Confocal images were acquired using a Leica SP5 confocal microscope with the pinhole set at 1 Airy unit. All images were processed with Fiji software. Images of cell bodies were acquired with z-stacks of 1 μm. To determine co-localisation, channels were viewed both individually and in composite. Co-localisation was determined if the cell body was visible in multiple channels through its entire thickness (multiple z planes). Representative examples of stacked images are shown. Images of axon terminals in nNOS+ neuron target areas were acquired with z-stacks of 0.5 μm. 10 Z-planes were stacked and brightness and contrast was adjusted equally across all axonal projection images for comparison. Images of synaptic terminals were acquired with z-stacks of 0.25 μm. ChR2-mCherry+ synaptic boutons were located in single z-planes, which were extracted from the stack to determine co-localisation with VGAT or VGluT2.

### Stereotaxic injections of AAV1-Ef1a-DIO-ChR2-mCherry

The DIO-ChR2-mCherry construct (gifted by the Deisseroth Lab) was commercially packaged in adeno-associated virus (AAV) serotype 2/1 vector consisting of the AAV2 ITR genomes and the AAV1 serotype capsid gene (Vector Biolab, Philadelphia). The viruses were diluted in sterile PBS and 5 % glycerol (pH 7.2) to a concentration of 2.7x10^13^ GC/ml. Unless otherwise stated, all viral tracing experiments were conducted on adult (11-13 weeks) NOS1Cre (-/+) mice. Mice were briefly anaesthetised in an induction chamber with isoflurane (4 %) and placed in a stereotaxic frame (David Kopf Instruments, California) with continued isoflurane administration (2 %). The eyes were protected with Lacri-lube, the scalp was shaved, and the skin disinfected with chlorheximide. All mice received a subcutaneous injection of carprofen (Rimadyl; 5 mg/kg) for post-operative anaesthesia. An incision (<1 cm) was made along the midline, and bupivacaine (2.5 mg/ml) was delivered directly to the incision site for local analgesia. A small hole was drilled in the scalp based on co-ordinates from bregma. Using a 33-gauge metal needle and a Hamilton syringe the virus solution (0.1 µl) was injected unilaterally at a flow rate of 0.3 µl/min. We systematically varied the injection co-ordinates (anterior-posterior; AP -3.0--3.4 mm, medial-lateral; ML 0.4-0.9 mm, dorsal-ventral; DV 4.3-4.8 mm) to obtain labelling of different sub-regions. The flow rate was controlled by a programmable pump (Elite Nanomite Infusion/Withdrawal Programmable Pump 11, 704507, Harvard Apparatus). After injection, the needle was left in place for 5 mins to allow for the spread of the virus. The incision was then sutured using nylon monofilament, non-absorbable sutures (size 2-0, 95060-062, VWR). Mice were allowed to recover in a heated chamber (30 °C) before being placed back into their home cage with littermates. All mice were monitored for five days after surgery, during which time they had access to carprofen (Rimadyl; 50 mg/ml) in their drinking water. Two weeks after surgery, the mice underwent transcardial perfusion, as described above, and tissue processed for microscopy.

### Experimental design & Statistics

#### Wild type Vs nNOS-deficient

To compare nNOS antibody staining in wild type and nNOS-deficient mice, the experimenter was blind to the strain of the mouse from the stage of immunolabelling until after image analysis. Mice for each experimental group were stained in parallel to control for differences between staining experiments. All images in this section were obtained with matched confocal settings. Each anti-nNOS antibody was tested in a total of three male WT and three male nNOS-deficient mice. The concentration of nNOS antibody was optimised through staining and imaging at three concentrations (1:250, 1:500, 1:1000). Images from the optimum concentration of 1:500 are shown.

#### Quantification of nNOS expressing neurons

For the quantification of nNOS expressing neurons triple immunolabelling for nNOS, HA, and TH was conducted in three male VGATCre:RiboTag mice. To obtain estimates of the numbers of nNOS neurons, and their neurotransmitter phenotype, every fourth midbrain section was selected for staining and imaging. Tilescans were taken of the entire VTA and SNc visible on the right-hand side of the brain section. Merged tilescan images were processed using Fiji (ImageJ) and VTA and SNc sub-region anatomy was defined based on TH expression. HA+ cells, nNOS+ cells and HA+/nNOS+ were counted in each sub-region using the ImageJ cell counter plugin.

#### nNOS neuron circuit tracing and ChR2-mCherry co-localisation

A total of 18 (8 males and 10 females) virus injected NOS1Cre mice exhibited ChR2-mCherry expression in the VTA and SNc. 8 of these mice also exhibited ChR2-mCherry expression in the supramamillary nucleus and were therefore excluded from further investigation. The remaining 10 mice were used to examine the axonal projections of nNOS+ neurons and further immunolabelling experiments. To investigate the co-localisation of ChR2-mCherry, nNOS and TH, images of sub-regions were processed using Fiji (ImageJ). All ChR2-mCherry+ cells were counted in each image using the ImageJ cell counter plugin.

#### Statistics

Data are presented as means + standard error of the mean (sem). Statistical comparisons were made using one-way ANOVA and Newman-Keuls post-hoc tests, where appropriate (Prism, Graphpad Software Inc).

## Results

### Comparison of three different anti-nNOS antibodies in the midbrain of wild type and nNOS-deficient mice

We first wanted to identify a reliable nNOS antibody for use in the VTA and SNc. We tested three different commercially available antibodies (see Table 1 & 2). We initially tested each antibody at three different concentrations (1:1000, 1:500, 1:250). For all three antibodies the 1:500 concentration appeared optimal in terms of reliably exhibiting immunolabelling in the interpeduncular nucleus (IPN) and in regions of the VTA and SNc in wildtype mice (Fig. 1). In order to thoroughly verify their specificity, each antibody (1:500) was used on midbrain sections from both wild type mice (*n* = 3) and nNOS-deficient mice (Huang *et al.*, 1993) (*n* = 3) as a negative control. It is well established that there is a large population of nNOS expressing neurons in the IPN, which lies just ventral to the VTA and was therefore well-suited to act as a positive control (Vincent & Kimura, 1992; Rodrigo *et al.*, 1994; Ascoli *et al.*, 2008). The first antibody (Sigma Aldrich (N7155; AB_260795)) failed to detect cell bodies and instead many processes were visible (Figure 1A), which were also present in the nNOS-deficient tissue, suggesting that it was non-specific. The second antibody (Cell Signalling (4234; AB_10694499)) displayed some sparse immunoreactivity ‘spots’ that could be mistaken for cell bodies within the VTA and SNc (Figure 1B), which were also present in the nNOS-deficient tissue, suggesting that they were non-specific. The third antibody (Sigma Aldrich (N2280; AB_260754)) exhibited clear immunolabelling of cell bodies in the wild-type tissue, which was completely absent in the nNOS-deficient tissue (Figure 1C). In the wildtype tissue nNOS+ neurons were mosaically distributed throughout the SNc, and most notably in the parabrachial pigmented nucleus (PBP) and rostral linear nucleus (RLi) of the VTA. These were in close proximity to TH+ neurons, but there is no co-localisation between nNOS and TH (Figure 1C).

**Figure 1.**
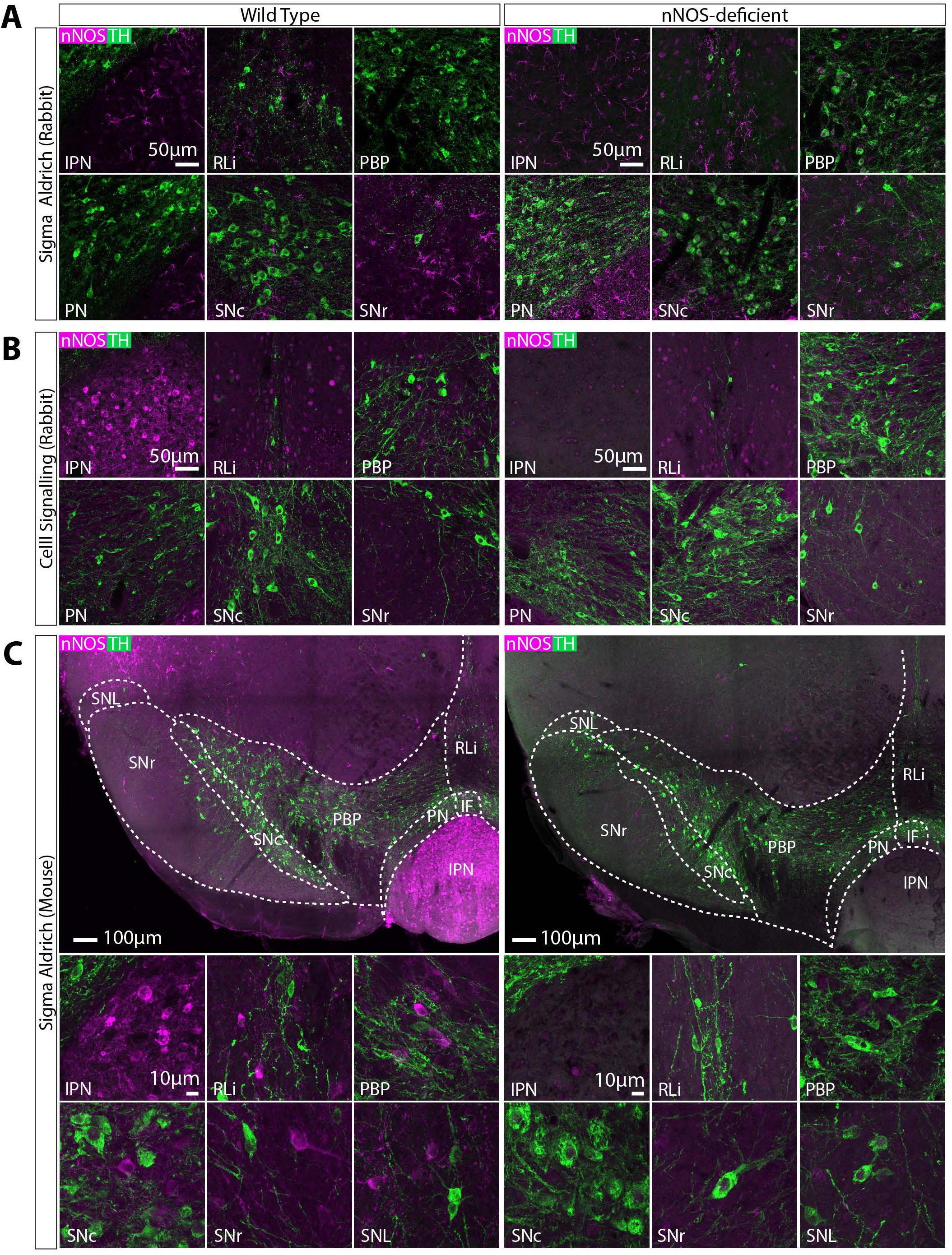
Comparison of three different anti-nNOS antibodies (see Table 1 & 2 for details) in the midbrain of wild type and nNOS-deficient mice. Representative images of double immunolabelling for nNOS (magenta) and TH (green). **A,** Anti-nNOS (Sigma Aldrich (N7155; AB_260795)) exhibited non-specific immunolabelling that was also seen in tissue from nNOS-deficient mice. **B**, Anti-nNOS (Cell Signalling (4234; AB_10694499)) also exhibited somewhat non-specific immunolabelling, that was only partially absent in tissue from nNOS-deficient mice. **C**, Anti-nNOS (Sigma Aldrich (N2280; AB_260754)) exhibited specific immunolabelling that was absent in tissue from nNOS-deficient mice. In wild type tissue, nNOS+ cells were observed in the IPN, RLi, PBP, SNc, SNr and SNL. There was no colocalisation between nNOS and TH.

**Table 2.** **Secondary antibodies**

| **Antibody** | **Conjugation** | **Host species** | **Supplier (cat#; RRID)** | **Concentration** |
| --- | --- | --- | --- | --- |
| Anti-Chicken | Alexa-488 | Goat | Thermo Fisher Scientific (A-11039; AB_2534096) | 1:1000 |
| Anti-Chicken | Cy3 | Donkey | Jackson ImmunoResearch Labs (703-165-155; AB_2340363) | 1:1000 |
| Anti-Mouse | Cy3 | Donkey | Jackson ImmunoResearch Labs (715-165-150; AB_2340813) | 1:1000 |
| Anti-Mouse | Cy5 | Donkey | Jackson ImmunoResearch Labs (715-175-151; AB_2340820) | 1:1000 |
| Anti-Rabbit | Alexa-633 | Goat | Thermo Fisher Scientific (A21070; AB_2535731) | 1:1000 |
| Anti-Rabbit | Cy3 | Donkey | Jackson ImmunoResearch Labs (711-165-152; AB_2307443) | 1:1000 |
| Anti-Goat | Alexa-488 | Donkey | Thermo Fisher Scientific (A11055; AB_2534102) | 1:1000 |
| Anti-Guinea Pig | Alexa-488 | Goat | Thermo Fisher Scientific (A11073; AB_2534117) | 1:1000 |

### nNOS is mostly expressed in GABAergic, non-dopaminergic neurons in the PBP part of the VTA and SNc, and mostly in non-GABAergic, non-dopaminergic (putatively glutamatergic) neurons in the VTAR and RLi

We next asked whether these nNOS+ neurons in the VTA and SNc were GABAergic. In the VTA and SNc, antibodies for markers of GABAergic identity (i.e., GABA, GAD and VGAT) do not robustly label cell bodies. We, therefore, used VGATCre mice (Vong *et al.*, 2011), where cre-recombinase is under the control of the promoter for VGAT, crossed with RiboTag mice (Sanz *et al.*, 2009) which contains a floxed hemagglutinin (HA)-tagged exon in the RLp22 gene. The resulting offspring (VGATCre:RiboTag) exhibit robust HA expression in cell bodies in the VTA and SNc which is well-suited to examining colocalisation using immunolabelling (somewhat more so than standard GFP and tdTomato reporter lines, in our hands). Triple immunolabelling for nNOS, HA, and TH was carried out in midbrain sections from VGATCre:RiboTag mice (*n* = 3 mice, 1420 neurons). Nuclei sub-regions were defined using TH immunolabelling and images from a mouse brain atlas (Franklin & Paxinos, 2008). All nNOS+ and HA+ neurons within each sub-region were counted. The number of nNOS+ neurons varied in different sub-regions with the largest populations lying in the PBP of the VTA and the RLi, with smaller populations found in the SNc and VTAR (ANOVA: F(3,8) = 22.33, p = 0.0003; Fig 2A-E). nNOS+ neurons were almost entirely absent in the interfascicular nucleus (IF) and the paranigral nucleus (PN) and therefore these sub-regions were not included in our analysis or further investigated.

**Figure 2.**
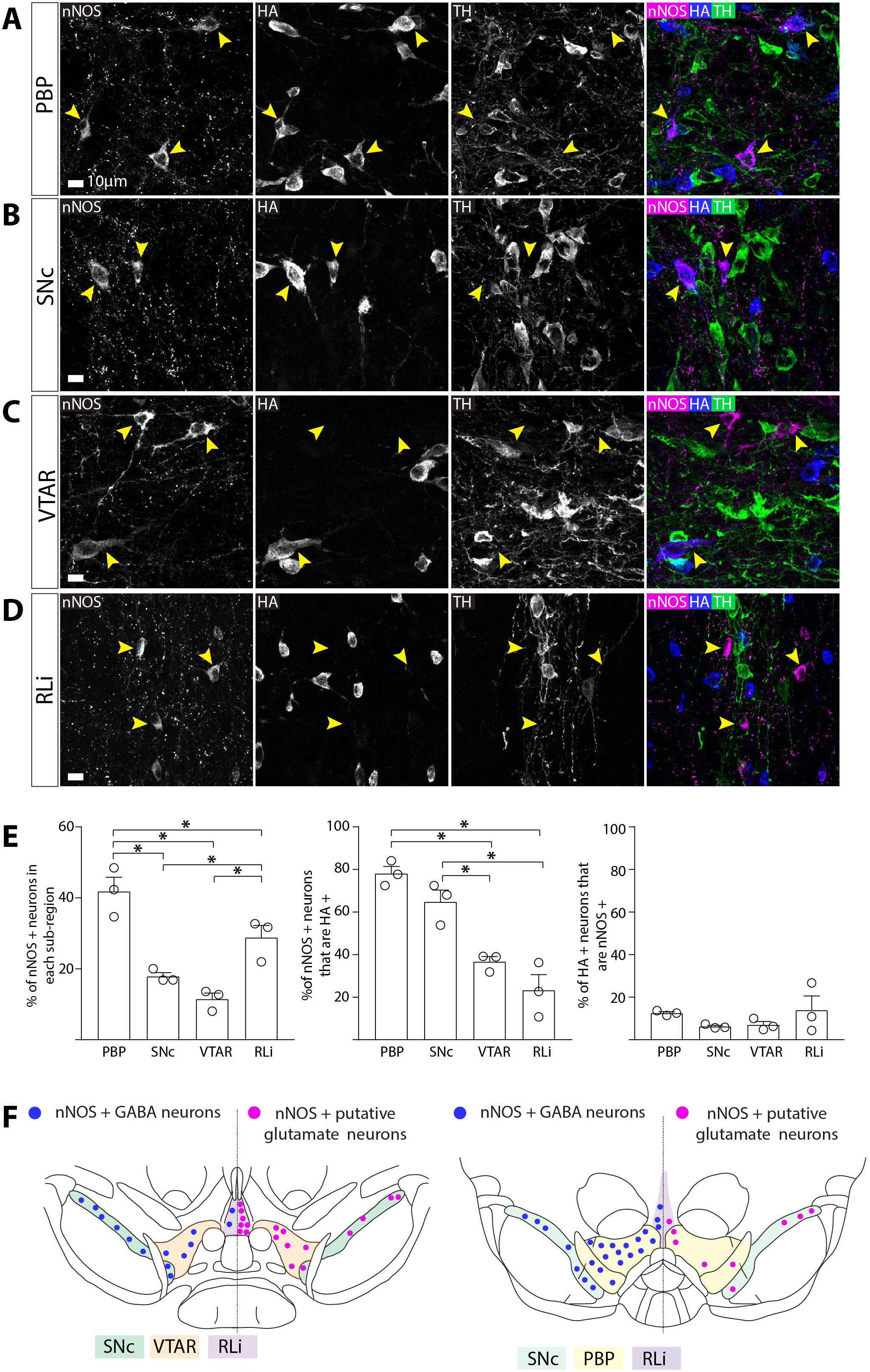
nNOS is mostly expressed in GABAergic, non-dopaminergic neurons in the PBP part of the VTA and SNc, and mostly in non-GABAergic, non-dopaminergic (putatively glutamatergic) neurons in the VTAR and RLi. **A-D,** Representative images of triple immunolabelling for nNOS, HA, and TH, in the SNc and sub-regions of the VTA. Yellow arrows indicate nNOS+ neurons. **E,** Graphs show the mean (+standard error of the mean; *n* = 3 mice, 1469 cells) and individual data points for percentage of nNOS+ cells localised in each region, percentage of nNOS+ cells that co-localised with HA, and the percentage of HA+ cells that were nNOS+. **F**, Schematic illustrating the localisation of nNOS+/GABAergic neurons and nNOS+/glutamatergic neurons in the VTA and SNc. Yellow arrows indicate exemplar neurons. * p < 0.05.

Consistent with our first set of results, there was no co-localisation between TH and nNOS. In contrast, co-localisation between nNOS and HA was extensive, although it varied between different sub-regions (ANOVA: F(3,8) = 24.54, p = 0.0002). In the PBP and SNc, the majority nNOS+ neurons were also HA+, suggesting that nNOS+ neurons in these regions are mostly GABAergic (Figure 2AB, EF). In contrast, in more rostral sub-regions (i.e., the VTAR and RLi) the majority of nNOS+ neurons were HA-- (and TH-) and therefore putatively glutamatergic (Figure 2CD, EF). Finally, the total proportion of HA+ neurons that expressed nNOS was similar in each sub-region (ANOVA: F(3,8) = 1.268, p = 0.3489; Figure 2E), typically less than 20%, indicating that nNOS+ neurons represent a sub-group of the overall GABAergic population in each of these subregions.

### AAV injection into the VTA and SNc of NOS1Cre+/-- mice lead to expression of ChR2-mCherry in cell bodies in distinct regions depending on injection volume/position

Having examined the neurochemical identity of nNOS+ neurons in the VTA and SNc, we next investigated their axonal projections. To do this, we did stereotaxic injections of AAV1-Ef1a-DIO-ChR2-mCherry into the midbrain of NOS1Cre+/-- mice (*n* = 18). We systematically varied the injection co-ordinates (see Methods) and then examined the degree of cell body expression of ChR2-mCherry within the SNc and VTA. We excluded mice that exhibited ChR2-mCherry expression in either the IPN or the SUM (both regions known to express nNOS; (Rodrigo *et al.*, 1994; Gonzalez-Hernandez & Rodriguez, 2000). The extent of cell body expression fell into three groupings (Figure 3; Table 3): Group 1 exhibited robust ChR2-mCherry cell body expression in the PBP, SNc, VTAR, and RLi; Group 2 exhibited robust ChR2-mCherry cell body expression in the PBP, SNc, and a dorso-lateral boundary region of the VTAR (where we did not see cell bodies in Group 1); Group 3 exhibited robust ChR2-mCherry cell body expression only in the PBP and SNc.

**Figure 3.**
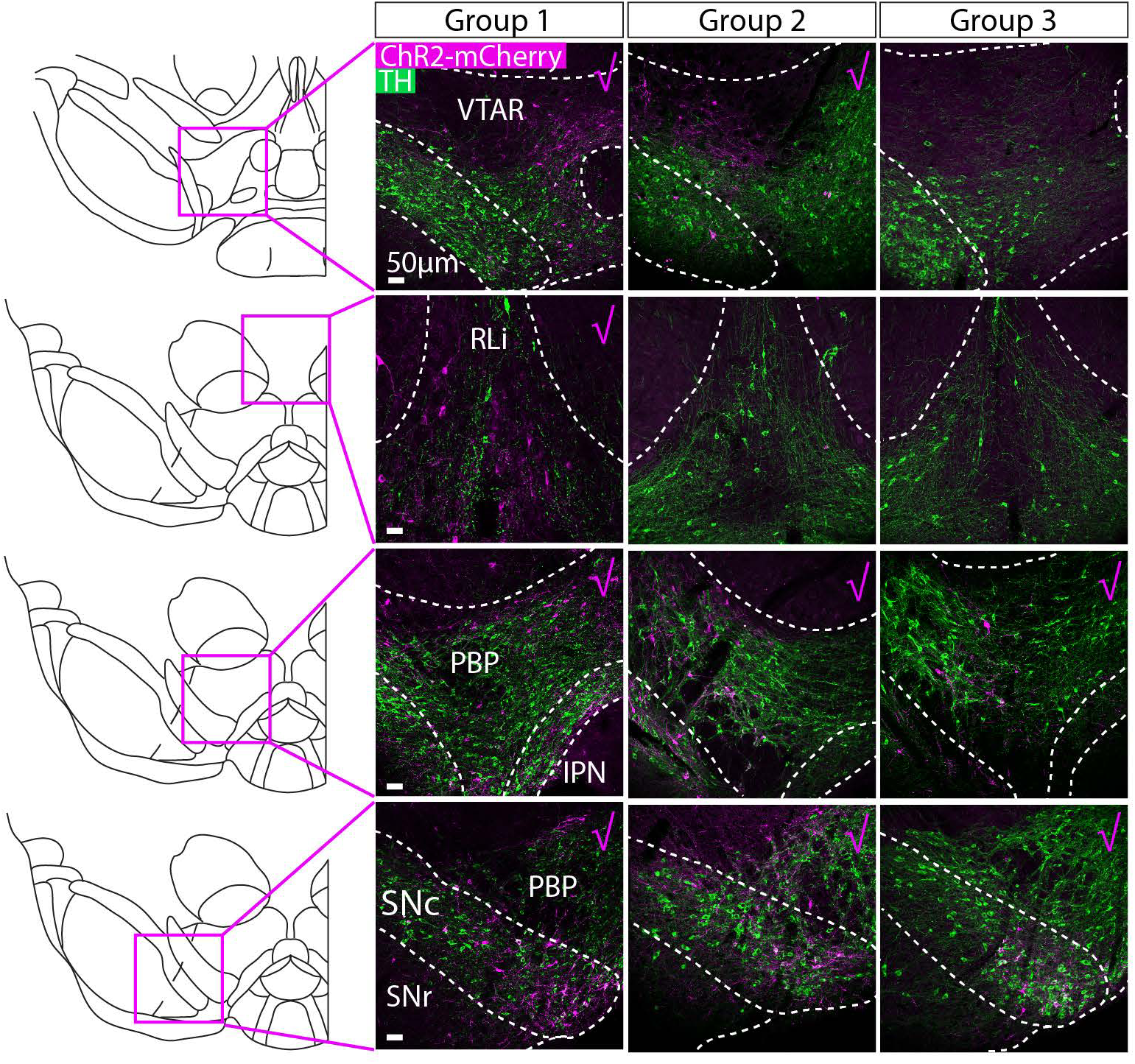
AAV injection into the VTA and SNc of NOS1cre+/-- mice lead to expression of ChR2-mCherry in cell bodies in distinct regions depending on injection volume/position. Representative images of ChR2-mCherry (magenta) and TH (green) in cell bodies for each injected group. Mice were grouped based on the distribution of ChR2-mCherry expression (pink tick indicates robust expression). Group 1 exhibited expression in the VTAR, RLi, PBP and SNc, Group 2 exhibited expression in the dorso-lateral VTAR, PBP, and SNc, and Group 3 exhibited expression that was restricted to the PBP and SNc (see Table 3).

**Table 3.** ChR2-mCherry expression density: +, very sparse expression; ++, modest expression; +++, dense expression. AA, Amygdaloid area; CLi, Caudal linear nucleus; DM, Dorsomedial hypothalamic nucleus; DR, Dorsal raphe nucleus; EAM, Extended amygdala, medial part; HDB, Horizontal limb of the diagonal band of Broca; LH, Lateral hypothalamus; LHb, Lateral habenula; LS, Lateral septum; MD, Dorsomedial nucleus of the hypothalamus; Mm, Mammillary bodies; MnR, Median raphe nucleus; MS, Medial septum; NAc, Nucleus accumbens; NRO, nucleus raphe obscurus PAG, Periaqueductal grey; PBP, Parabrachial pigmented nucleus; PMnR, Paramedian raphe nucleus; PO, Preoptic area; RLi, Rostral linear nucleus; SNr, Substantia nigra pars reticulata; ST, stria terminalis; VM, Ventromedial thalamus; VP, ventral pallidum; VTAR, Rostral ventral tegmental area; ZI, Zona inserta. ^a^, cell bodies were restricted to the dorso-lateral boundary region of the VTAR and this was not seen in Group 1 or Group 3.

|  | **Group 1** |  |  |  | **Group 2** |  |  | **Group 3** |  |  |
| --- | --- | --- | --- | --- | --- | --- | --- | --- | --- | --- |
| **Injection no.** | **7** | **8** | **9** | **10** | **13** | **14** | **15** | **16** | **17** | **18** |
| **Cell Bodies** |  |  |  |  |  |  |  |  |  |  |
| VTAR | + | ++ | ++ | +++ | +++ | +++ | +++ |  | + | + |
| RLi | +++ | +++ | +++ | +++ |  |  |  |  |  |  |
| PBP | +++ | +++ | +++ | +++ | +++ | +++ | +++ | ++ | +++ | ++ |
| SNc | +++ | +++ | +++ | +++ | +++ | +++ | +++ | ++ | +++ | + |
| **Axonal Projections** |  |  |  |  |  |  |  |  |  |  |
| **Septum** |  |  |  |  |  |  |  |  |  |  |
| MS | ++ | ++ | ++ | ++ |  |  |  |  |  |  |
| HDB | + | + | + | + |  |  |  |  |  |  |
| LS | + | + | + | + |  |  |  |  |  |  |
| ST | + | + | + | + |  |  |  |  |  |  |
| **Striatum** |  |  |  |  |  |  |  |  |  |  |
| NAc (shell) | + | + | + | + |  |  |  |  |  |  |
| VP | +++ | +++ | +++ | +++ |  |  |  |  |  |  |
| NAc (core) |  |  |  | + |  |  |  |  |  |  |
| **Hypothalamus** |  |  |  |  |  |  |  |  |  |  |
| LH | +++ | +++ | +++ | +++ | + | + | + |  |  |  |
| DM | + | + | + | + |  |  | + |  |  |  |
| PO | ++ | ++ | ++ | ++ |  |  |  |  |  |  |
| ZI | + | + | + | + |  |  |  |  |  |  |
| **Thalamus** |  |  |  |  |  |  |  |  |  |  |
| LHb | + | + | + | + |  |  |  |  |  |  |
| MD |  |  |  | + |  |  |  |  |  |  |
| VM | + | + | + | + |  |  |  |  |  |  |
| **Amygdala** |  |  |  |  |  |  |  |  |  |  |
| EAM |  |  |  | + |  |  |  |  |  |  |
| AA |  |  |  | + |  |  |  |  |  |  |
| **Midbrain** |  |  |  |  |  |  |  |  |  |  |
| Mammillary | + |  |  |  |  |  |  |  |  |  |
| **Pons/medulla** |  |  |  |  |  |  |  |  |  |  |
| DR | + | + | + | ++ |  |  |  |  |  |  |
| PMnR | + | + | + | ++ |  |  |  |  |  |  |
| MnR | ++ | ++ | + | ++ |  |  |  |  |  |  |
| PAG | + | + | + | + |  |  |  |  |  |  |
| CLi | + | + | + | + |  |  |  |  |  |  |
| NRO | + | + | + | + |  |  |  |  |  |  |

### When ChR2-mCherry expression was restricted to cell bodies in the PBP part of the VTA and the SNc, no axonal projections were found outside of the VTA and SNc

For each mouse we conducted a full survey of the entire brain looking for ChR2-mCherry positive axonal projections. In brains from Group 1 (which exhibited cell body labelling in the PBP, SNc, VTAR, and RLi), we observed extensive axonal projections multiple regions (Figure 4; Table 3), all shown previously to receive input from GABA and glutamate neurons in the VTA (Taylor *et al.*, 2014). These projections were most dense in the ventral pallidum (VP), lateral hypothalamus (LH), and median raphe (MnR). In brains from Group 2 (which exhibited cell body in the PBP, SNc and dorso-lateral part of the VTAR) we only reliably observed very sparse processes in the LH (Figure 4; Table 3). In brains from Group 3 (which exhibited robust cell body labelling only in the PBP and SNc) we did not observe any axonal projections outside of the VTA and SNc (Figure 4; Table 3). On the basis of these expression patterns we can, therefore, draw two main conclusions. First, NOS1Cre+ neurons in the PBP and SNc do not send axonal projections outside of the VTA and SNC. Second, NOS1Cre+ neurons in the VTAR and RLi send extensive projections to multiple regions, including the VP, LH and MnR. All of these regions are known to receive input from the RLi (Del-Fava *et al.*, 2007). It should be noted that in the case of Group 2, where some sparse fibres were observed the LH, the cell body labelling in these cases was restricted to the dorso-lateral part of the VTAR only. In contrast in Group 1 cell body labelling was observed throughout the VTAR.

**Figure 4.**
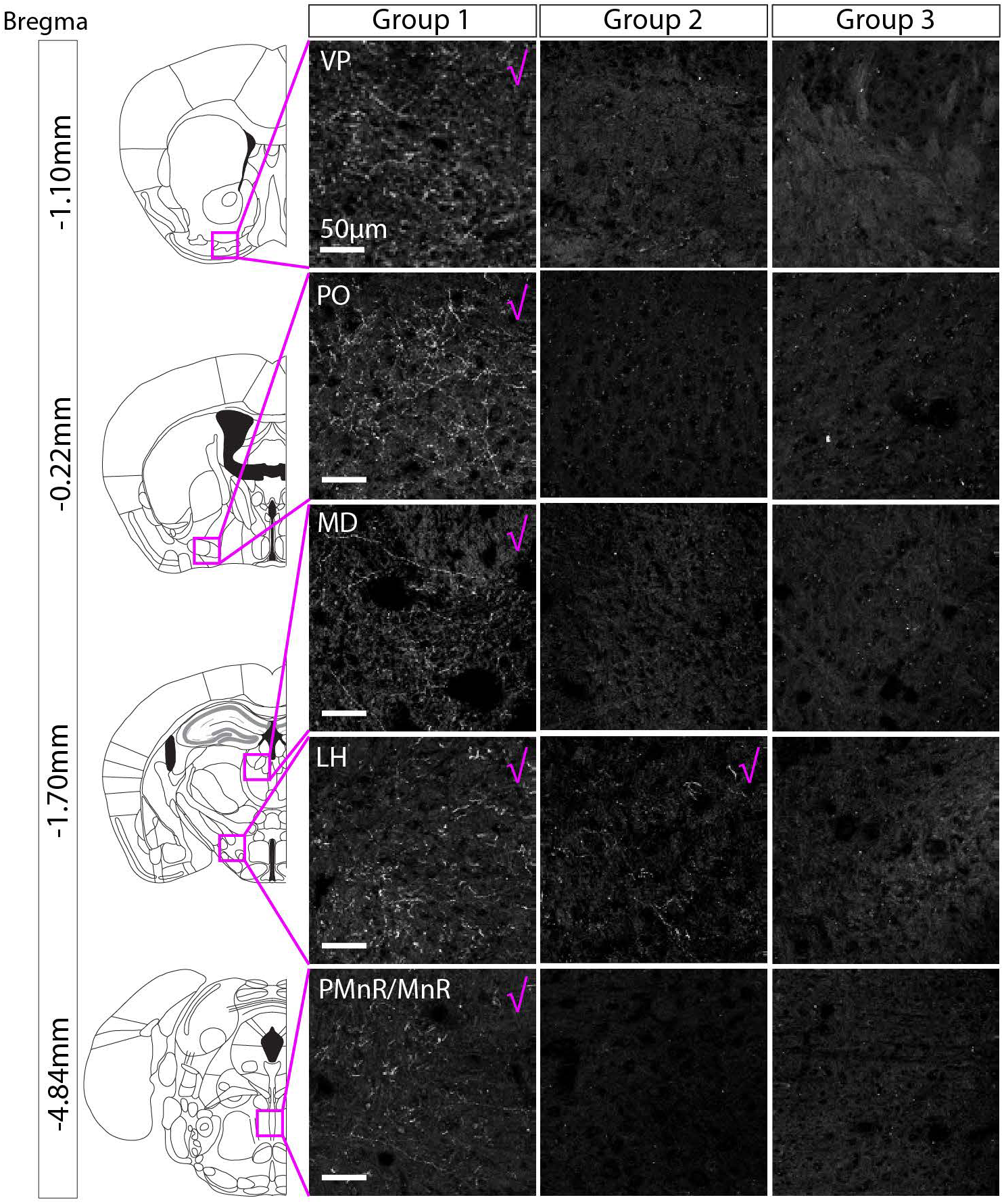
When ChR2-mCherry expression was restricted to cell bodies in the PBP part of the VTA and the SNc, no axonal projections were found outside of the VTA and SNc. Representative images of axon-expressed ChR2-mCherry for each group. Group 1 exhibited extensive projections (see Table 3 for full summary) to multiple regions. Images are shown for the VP, PO, MD, LH, IPN, and PMnR/MnR, where the most extensive axonal expression was observed (pink tick indicates robust axonal expression). Group 2 exhibited sparse projections that were limited to the LH. Group 3 (which had cell body labelling restricted to the PBP and SNc) did not exhibit any axonal expression outside of the VTA and SNc.

### Cell body expression of ChR2-mCherry was colocalised with nNOS immunolabelling in the VTA, but in the SNc some neurons were TH+

We next examined the degree of colocalisation between ChR2-mCherry, nNOS, and TH in cell bodies in the PBP, SNc, VTAR and RLi (n = 3-5 mice, 554 neurons). We conducted immunolabelling for nNOS and TH and examined co-localisation with ChR2-mCherry. In the PBP (*n* = 5 mice; ANOVA: F(2, 12) = 290.0, p < 0.0001), VTAR (*n* = 4 mice; ANOVA: F(2,9) = 35.27, p < 0.0001), and RLi (*n* = 3 mice; ANOVA: F(2,6) = 213.9, p < 0.0001) nucleus, almost all ChR2-mCherry+ cells were nNOS+ and TH-- (Figure 5A-C).

**Figure 5.**
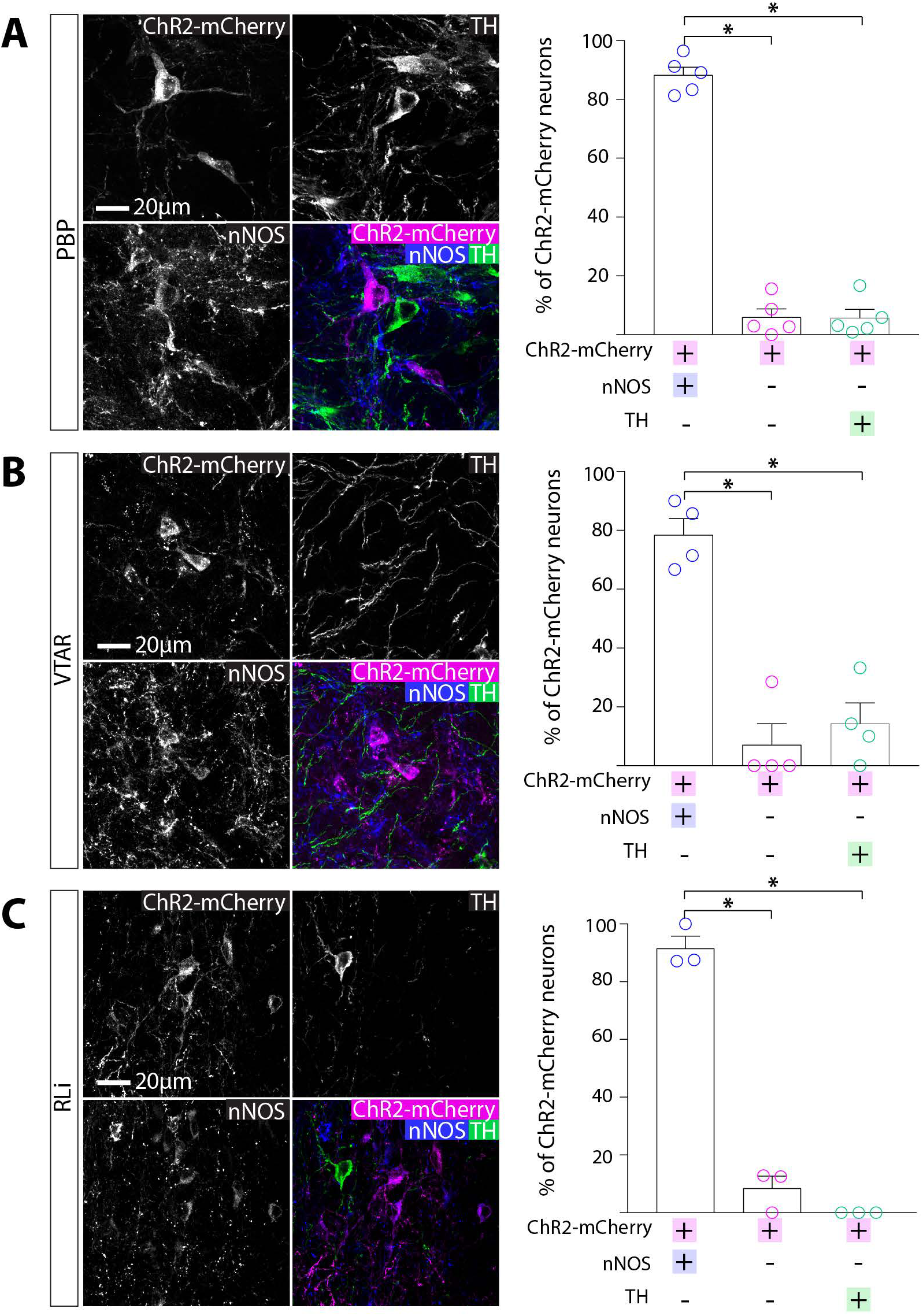
Cell body expression of ChR2-mCherry was colocalised with nNOS immunolabelling in the VTA. **A,** Representative images of triple immunolabelling for ChR2-mCherry, nNOS, TH in the PBP. Graph shows the mean (+standard error of the mean) and individual data points for percentage of ChR2-mCherry that were co-localised with nNOS and/or TH (230 ChR2-mCherry+ cells, 5 mice). Almost all ChR2-mCherry+ neurons were nNOS+ and TH-. **B,** Representative images of triple immunolabelling for ChR2-mCherry, nNOS, TH in the VTAR. Graph shows the mean (+standard error of the mean) and individual data points for percentage of ChR2-mCherry that were co-localised with nNOS and/or TH (40 ChR2-mCherry+ cells, 4 mice). Almost all ChR2-mCherry+ neurons were nNOS+ and TH-. **C,** Representative images of triple immunolabelling for ChR2-mCherry, nNOS, TH in the RLi part of the VTA. Graph shows the mean (+standard error of the mean) and individual data points for percentage of ChR2-mCherry that were co-localised with nNOS and/or TH (155 ChR2-mCherry+ cells, 3 mice). Almost all ChR2-mCherry+ neurons were nNOS+ and TH-. * p < 0.05.

In contrast, in the SNc similar numbers of neurons were ChR2-mCherry+ and/or nNOS+ and/or TH+ (ANOVA: F(2,12) = 2.627, p = 0.1132). Although a majority of the ChR2-mCherry+ cells were nNOS+ (Figure 6), surprisingly, around half of the ChR2-mCherry+ neurons in the SNc were TH+ (and nNOS--; Figure 6). As observed in both the wild-type and VGATCre:RiboTag mice, nNOS antibody immunolabelling did not co-localise with TH in the SNc. It is possible, however, that these neurons appear nNOS-- because they are either expressing very low levels of the enzyme (so that it is not detectable with the nNOS antibody), or that nNOS mRNA is being transcribed but the protein is not being synthesised currently.

**Figure 6.**
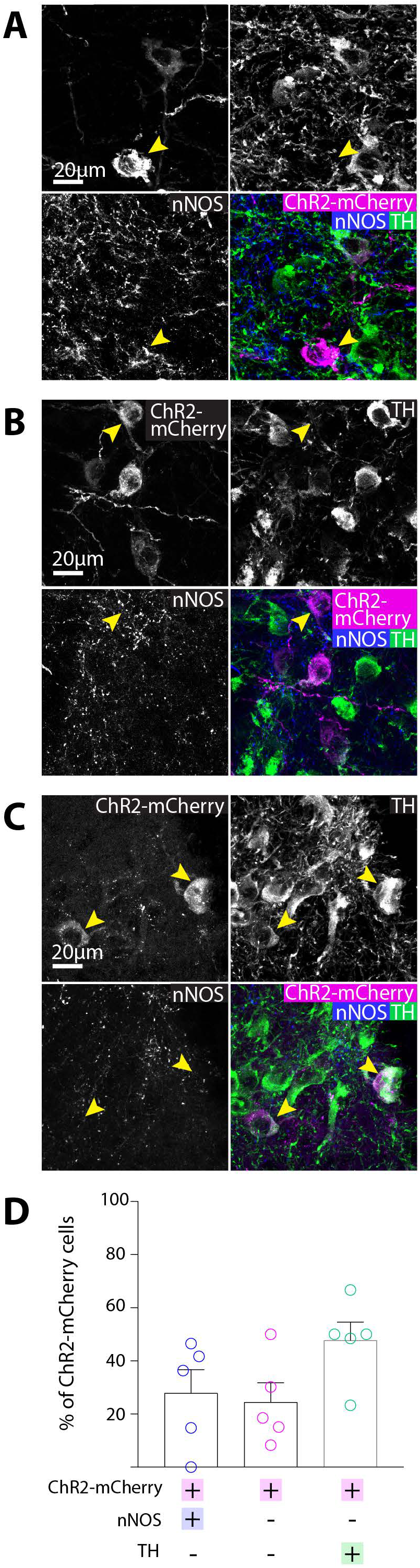
Cell body expression of ChR2-mCherry was mostly colocalised with either nNOS or TH in the SNc. **A,** Representative images of triple immunolabelling for ChR2-mCherry, nNOS, TH in the SNc, showing co-localisation of nNOS and ChR2-mCherry. **B,** Representative images of triple immunolabelling for ChR2-mCherry, nNOS, TH in the SNc, showing colocalisation of TH and ChR2-mCherry. **C**, Graph shows the mean (+standard error of the mean) and individual data points for percentage of ChR2-mCherry that were co-localised with nNOS and/or TH (129 ChR2-mCherry+ cells, 5 mice). Most ChR2-mCherry+ neurons were either nNOS+ or TH+. Yellow arrows indicate exemplar neurons.

### Axonal expression of ChR2-mCherry+ was colocalised with GABAergic synaptic boutons in the VTA and SNc

Taken together, our findings suggest that nNOS+ neurons in the PBP and SNc are GABAergic and do not project outside the VTA and SNc. To further examine their neurochemical identity, we examined single-z-plane images of tissue immunolabelled for VGAT and TH. In the VTA and SNc, although VGAT antibodies do not resolve cell bodies well (as discussed earlier) they can reliably label processes, include putative presynaptic boutons. In the VTA, we commonly observed VGAT+ puncta co-localised with ChR2-mCherry and in close proximity to, but not co-localising with, TH+ processes (Figure 7A). This is consistent with the possibility that nNOS+ interneurons form inhibitory synapses onto dopamine neurons. In addition, in the SNc we were able to locate some ChR2-mCherry+ fibres that were also colocalised with VGAT+/TH+ puncta (Figure 7B).

**Figure 7.**
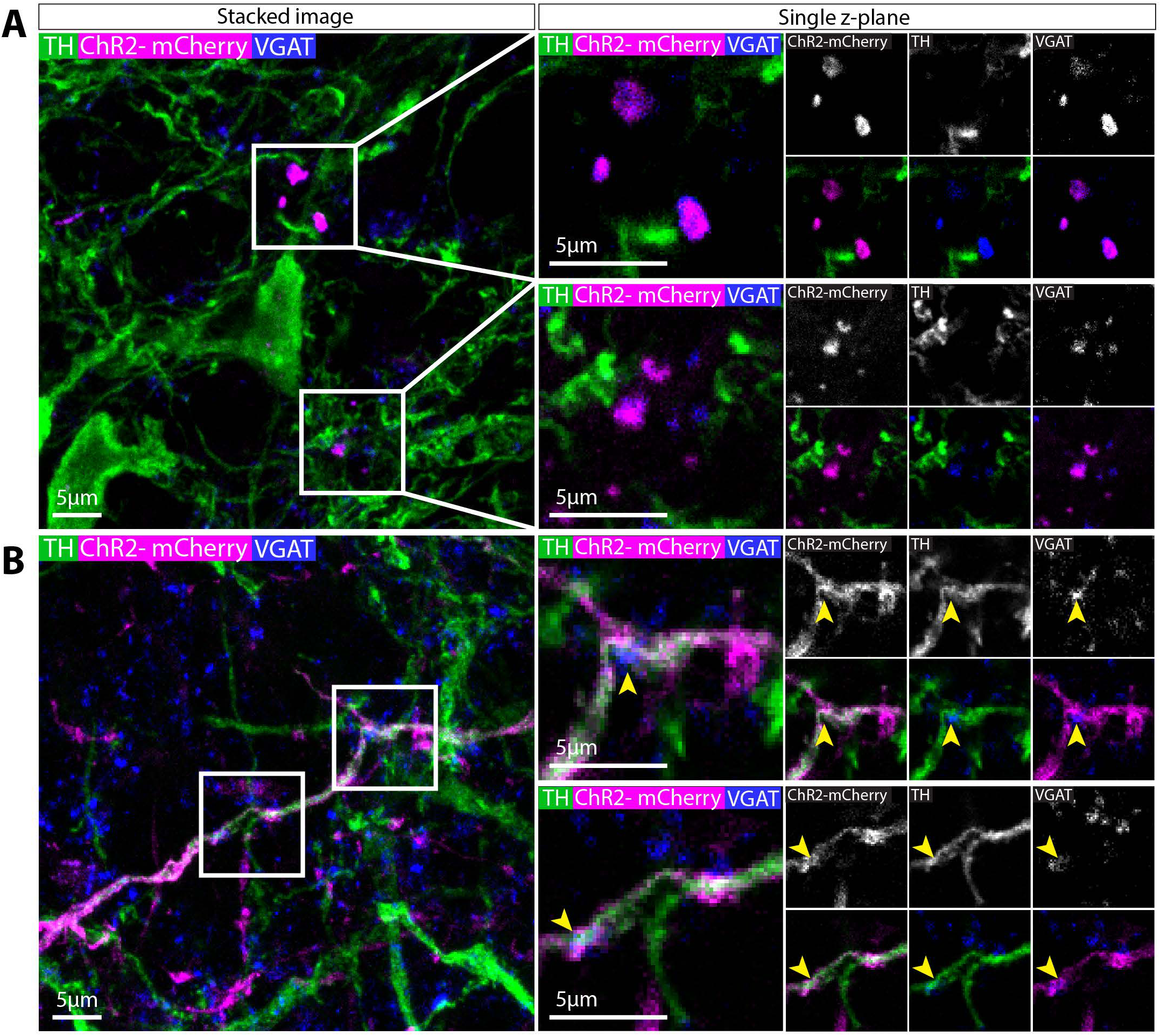
Axonal expression of ChR2-mCherry+ was colocalised with GABAergic synaptic boutons in the VTA and SNc. **A,** Representative images of immunolabelling for ChR2-mCherry, VGAT, and TH in the PBP. ChR2-mCherry co-localises with VGAT puncta in a single z-plane image suggesting the presence of GABAergic synapses. **B**, Representative images of immunolabelling for ChR2-mCherry, VGAT, and TH in the SNc. ChR2-mCherry co-localises with VGAT puncta in a single z-plane image suggesting the presence of GABAergic synapses. These puncta are also often TH+. Yellow arrows indicate exemplar puncta.

### Axonal expression of ChR2-mCherry was colocalised with glutamatergic synaptic boutons in the VP and MnR

The VP is the area that received the most prominent input from the nNOS+ neurons in the RLi nucleus, consistent with non-cell-type-specific anterograde tracing approaches (Del-Fava *et al.*, 2007). To examine this innervation in more detail, VP containing sections were immunolabelled for substance P (which delineates the VP) and either VGluT2 or VGAT. It can be clearly seen that ChR2-mCherry+ fibres were more prevalent in the VP (substance P+ region) compared to the horizontal limb of the diagonal band of Broca (HDB) and shell of the NAc (areas that receive sparse innervation; Figure 8A). This innervation is present throughout the extent of the VP. ChR2-mCherry+ puncta could be clearly visualised amongst substance P+ puncta, and were commonly colocalised with VGluT2+ puncta (Figure 8B). This is consistent with our observation that these projections originate mostly from cell bodies in the RLi and VTAR that are VGAT-/TH-- and therefore putatively glutamatergic. Indeed, when we examined VGluT2 and ChR2-mCherry colocalisation in the RLi, we observed some VGluT2+ cell bodies (as for GABAergic markers, it can be difficult to resolve cell bodies with antibodies for markers of glutamatergic neurons in the VTA) that were ChR2-mCherry+, consistent with our hypothesis that this is a predominantly glutamatergic population (Figure 8C). Lastly when we conducted immunolabelling for VGAT we occasionally observed colocalisation with ChR2-mCherry+ puncta, but these were less common than for VGluT2 (Figure 8D).

**Figure 8.**
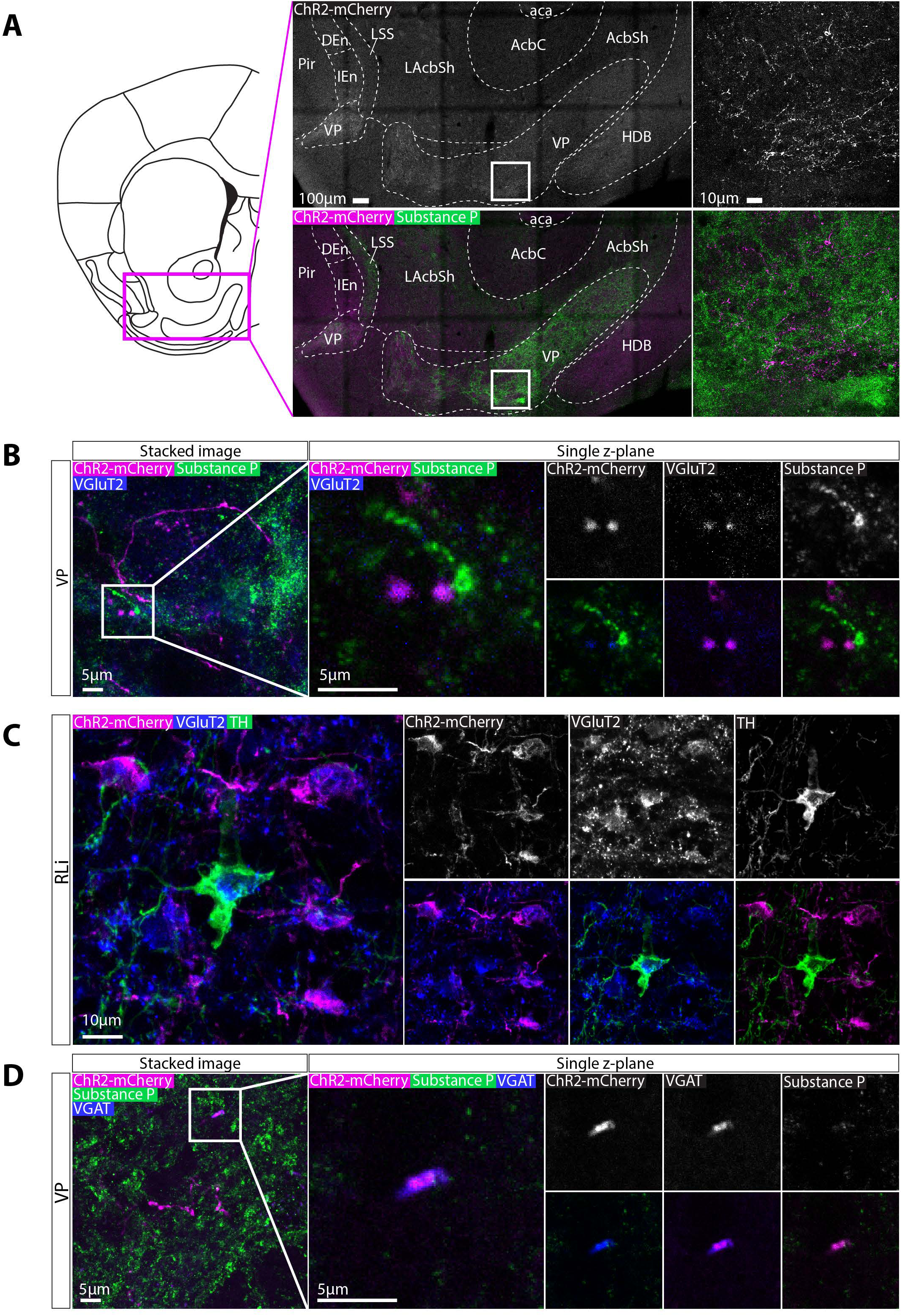
Axonal expression of ChR2-mCherry was colocalised with glutamatergic synaptic boutons in the VP. **A,** Representative images of immunolabelling for ChR2-mCherry and Substance P (which is highly expressed in the VP). Extensive innervation was observed in the VP compared to the neighbouring parts of the NAc and septum. **B**, High-magnification representative images of immunolabelling for ChR2-mCherry, Substance P, and VGluT2 in the VP. Colocalisation between ChR2-mCherry andVGluT2 puncta can be seen in single zplane images, suggesting that these projections are glutamatergic. **C,** Representative images of immunolabelling for ChR2-mCherry, VGluT2, and TH, in the RLi, occasionally also revealed cell bodies that expressed VGluT2. **D,** High-magnification representative images of immunolabelling for ChR2-mCherry, Substance P, and VGAT in the VP. On some occasions, colocalisation between ChR2-mCherry and VGAT puncta was observed in single z-plane images, suggesting that some these projections are also be GABAergic.

A second region that received extensive input was the MnR. Immunolabelling for serotonin (5-HT) revealed ChR2-mCherry+ terminals often in close proximity 5-HT+ neurons. (Figure 9A). Similar to the VP, VGluT2+ (Figure 9B) and VGAT+ (Figure 9C) puncta co-localised with ChR2-mCherry+ puncta in single z-plane images.

**Figure 9.**
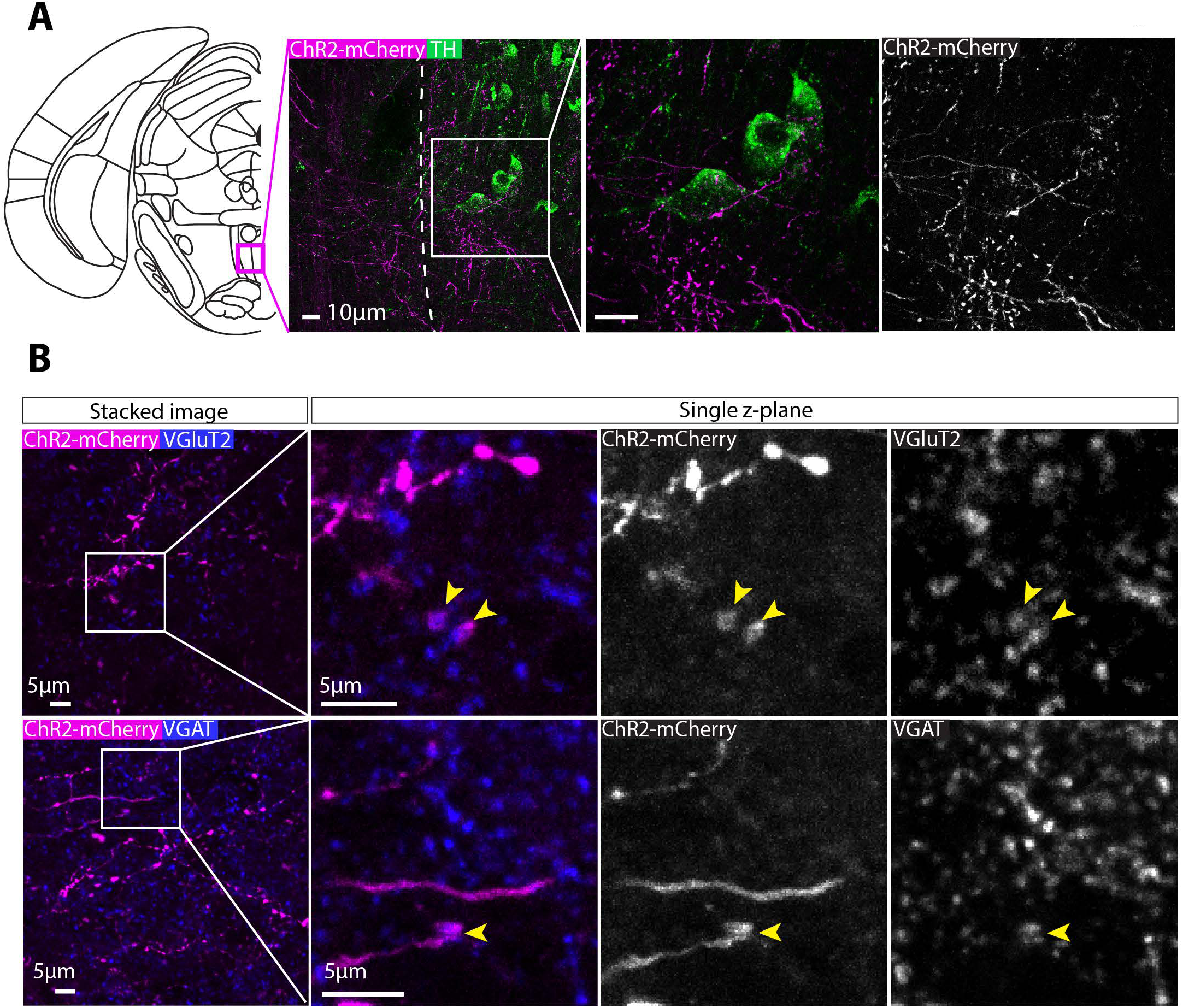
Axonal expression of ChR2-mCherry was colocalised with glutamatergic synaptic boutons in the MnR. **A,** Representative images of immunolabelling for ChR2-mCherry and 5-HT (which is highly expressed in the MnR compared to nearby regions). Extensive innervation was observed in the MnR with axons often passing in close apposition to 5-HT+ neurons. **B**, High-magnification representative images of immunolabelling for ChR2-mCherry and VGluT2 in the MnR. Colocalisation between ChR2-mCherry and VGluT2 puncta can be seen in single z-plane images, suggesting that these projections are glutamatergic. **C,** High-magnification representative images of immunolabelling for ChR2-mCherry and VGAT puncta in the VP. On some occasions, colocalisation between ChR2-mCherry and VGAT was observed in single z-plane images, suggesting that some these projections may also be GABAergic.

## Discussion

Previous investigations of nNOS expression in the VTA and SNc have produced discrepant results with respect to the extent of its expression, which subregions within the VTA and SNc it is expressed in, and the degree of co-expression by dopamine neurons (Vincent & Kimura, 1992; Rodrigo *et al.*, 1994; Gonzalez-Hernandez & Rodriguez, 2000; Klejbor *et al.*, 2004; Gotti *et al.*, 2005; Cavalcanti-Kwiatkoski *et al.*, 2010; Mitkovski *et al.*, 2012). We hypothesised that this variation in the literature was in part due to the use of different antibodies not validated specifically for the VTA and SNc. Consistent with this we found that 3 antibodies tested produced divergent results and in only one case was all immunolabelling absent in control tissue from nNOS-deficient mice. This highlights the importance of validating antibodies. Using our validated antibody we show that nNOS+ neurons are present in the SNc, VTAR, PBP, and RLi, but not other parts of the VTA including the PN. In addition, we show that nNOS+ neurons in the SNc and PBP are largely GABAergic, whereas those located in the RLi and VTAR are largely glutamatergic. These GABAergic neurons appear to be interneurons: despite the high levels of expression of an anterograde tracer in their cell bodies, we could not detect any axonal projections outside of the VTA and SNc. Across these regions nNOS+ neurons interneurons make up less than 10% of the total GABA neuron population. In contrast, we found that nNOS+/glutamatergic neurons sent extensive projections to several regions, including the VP, LH and MnR.

Previously it has been demonstrated that GABA neurons in the VTA make anatomically-defined local synaptic connections with dopamine and non-dopamine neurons in the VTA (Omelchenko & Sesack, 2009). Moreover, functional optogenetic stimulation of VTA GABA neurons can evoke fast GABAA-receptor-mediated synaptic currents in dopamine neurons in the VTA (Tan *et al.*, 2012; van Zessen *et al.*, 2012). Activation of this local GABAergic microcircuit can generate a conditioned place aversion and reduce food consumption (Tan *et al.*, 2012; van Zessen *et al.*, 2012). It was not clear, however, whether the GABA neurons that made these local synaptic connections were also the same GABA neurons that send long-range projections to other regions such as the striatum (Brown *et al.*, 2012; Taylor *et al.*, 2014). Our findings indicate that at least one subset of these neurons are local GABAergic interneurons. Moreover, because these neurons have a distinct molecular identity (i.e., nNOS expression) they are experimentally tractable (e.g., by using cell-type-specific functional and anatomical techniques in NOS1Cre mice). This approach could be further refined using intersectional genetics (e.g., to limit expression based GABAergic or glutamatergic identity).

One intriguing observation was that a subset of nNOS-Cre+ neurons in the SNc were TH+. Importantly, these neurons do not send projections outside the SNc. It is, of course, a canonical view of the mesocorticolimbic dopamine system that TH+ neurons in the SNc send extremely dense axonal projections to several target regions, most notably the striatum (Matsuda *et al.*, 2009). Our findings suggest, however, that a subset of TH+ neurons in the SNc are local interneurons. Interestingly, there is evidence for TH+ GABAergic interneurons in the striatum (Dubach *et al.*, 1987; Tashiro *et al.*, 1989; Meredith *et al.*, 1999; O’Byrne *et al.*, 2000; Mao *et al.*, 2001; Petroske *et al.*, 2001; Ibanez-Sandoval *et al.*, 2010; Unal *et al.*, 2011; Unal *et al.*, 2015; Xenias *et al.*, 2015). Optogenetic stimulation of these TH+ neurons in the striatum fails to elicit any detectable release of dopamine (Ibanez-Sandoval *et al.*, 2010; Xenias *et al.*, 2015). In addition, they do not express AADC, dopamine, or DAT (Xenias *et al.*, 2015). Instead, optogentic activation of these neurons elicited GABA-mediated IPSCs in mediam spiny neurons (Ibanez-Sandoval *et al.*, 2010; Xenias *et al.*, 2015). Co-localistation with GABA synthesising enzymes GAD65 and GAD67 has also been reported (Betarbet *et al.*, 1997; Cossette *et al.*, 2005; Mazloom & Smith, 2006; Tandé *et al.*, 2006; San Sebastián *et al.*, 2007). Notably, this interneuron population is considered to be distinct from the nNOS+ interneurons in the striatum (Ibanez-Sandoval *et al.*, 2010; Tepper *et al.*, 2010b). It will be important, therefore, to establish if TH+ interneurons in the SNc are dopaminergic. Our examination of synaptic terminals in the SNc suggest that some at least may be GABAergic. It is also not clear if these TH+ interneurons would be mistaken for TH+, long-range projecting dopamine neurons in studies where TH-GFP or TH-Cre mice are used to identify and/or manipulate dopamine neurons.

Glutamate neurons are found sparsely distributed throughout the SNc and VTA, although at a greater density in more medial regions of the VTA (Yamaguchi *et al.*, 2007; Nair-Roberts *et al.*, 2008; Yamaguchi *et al.*, 2011; Yamaguchi *et al.*, 2013; Morales & Root, 2014; Yamaguchi *et al.*, 2015; Root *et al.*, 2016; Morales & Margolis, 2017). Some of these neurons co-release dopamine or GABA (Stuber *et al.*, 2010; Tecuapetla *et al.*, 2010; Root *et al.*, 2014b; Zhang *et al.*, 2015; Yoo *et al.*, 2016). They make local synaptic connections with dopamine and non-dopamine neurons and send projections to several regions including the striatum (Dobi *et al.*, 2010; Hnasko *et al.*, 2012; Root *et al.*, 2014a; Root *et al.*, 2014b; Taylor *et al.*, 2014). Interestingly, optogenetic excitation of VTA glutamate neurons can have rewarding and aversive effects, depending in part on the site of stimulation, suggesting some functional heterogeneity (Root *et al.*, 2014a; Wang *et al.*, 2015; Qi *et al.*, 2016; Yoo *et al.*, 2016). We have found the nNOS is expressed by glutamate neurons in the VTAR and RLi that send projections most densely to the VP, LH, and MnR. This is consistent with previous reports of non-cell-type specific anterograde labelling of projections from the RLi to the VP, but not NAc (Del-Fava *et al.*, 2007). Based on reports of the full projectome of glutamate neurons in the VTA, which includes extensive projections to regions such as the NAc, (Hnasko *et al.*, 2012; Taylor *et al.*, 2014; Qi *et al.*, 2016), we conclude that nNOS+ neurons represent a projection-specific sub-group of this population. As is the case for nNOS+ GABA neurons in the PBP and SNc, because nNOS+ glutamate neurons have a distinct molecular identity understanding their function will be experimentally tractable.

In conclusion, our findings indicate that nNOS is expressed by neurochemically-- and anatomically-distinct neuronal sub-groups in a sub-region-specific manner within the VTA and SNc.

## Acknowledgements

This work was supported by grant**s** MC-A654-5QB70 (M.A.U.), MCA654-5QB60 (J.L.), and MC-A654-5QB40 (D.J.W.) from the U.K. Medical Research Council (MRC).

## References

Ascoli GA et al (2008) Petilla terminology: nomenclature of features of GABAergic interneurons of the cerebral cortex. Nat Rev Neurosci 9: 557–568.

Backes E, Hemby SE (2003) Discrete cell gene profiling of ventral tegmental dopamine neurons after acute and chronic cocaine self-administration. J Pharmacol Exp Ther 307: 450–459.

Betarbet R, Turner. R, Chockkan V, DeLong MR, Allers KA, Walters J, Levey AI, Greenamyre JT (1997) Dopaminergic neurons intrinsic to the primate striatum. J Neurosci 17: 6761–6768.

Brown MT, Tan KR, O’Connor EC, Nikonenko I, Muller D, Luscher C (2012) Ventral tegmental area GABA projections pause accumbal cholinergic interneurons to enhance associative learning. Nature 492: 452–456.

Cavalcanti-Kwiatkoski R, Raisman-Vozari R, Ginestet L, Del Bel E (2010) Altered expression of neuronal nitric oxide synthase in weaver mutant mice. Brain Res 1326: 40–50.

Cossette M, Lecomte F, Parent A (2005) Morphology and distribution of dopaminergic neurons intrinsic to the human striatum. J Chem Neuroanat 29: 1–11.

Del-Fava F, Hasue RH, Ferreira JG, Shammah-Lagnado SJ (2007) Efferent connections of the rostral linear nucleus of the ventral tegmental area in the rat. Neuroscience 145: 1059–1076.

Dobi A, Margolis EB, Wang HL, Harvey BK, Morales M (2010) Glutamatergic and nonglutamatergic neurons of the ventral tegmental area establish local synaptic contacts with dopaminergic and nondopaminergic neurons. J Neurosci 30: 218–229.

Dougalis AG, Matthews GA, Bishop MW, Brischoux F, Kobayashi K, Ungless MA (2012) Functional properties of dopamine neurons and co-expression of vasoactive intestinal polypeptide in the dorsal raphe nucleus and ventro-lateral periaqueductal grey. Eur J Neurosci 36: 3322–3332.

Dubach M, Schmidt R, Kunkel D, Bowden DM, Martin R, German DC (1987) Primate neostriatal neurons containing tyrosine hydroxylase: immunohistochemical evidence. Neurosci Lett 75: 205–210.

Franklin KBJ, Paxinos G (2008) The mouse brain in stereotaxic coordinates. Acad. Press, [Amsterdam] [u.a.

Garthwaite J (1991) Glutamate, nitric oxide and cell-cell signalling in the nervous system. Trends Neurosci 14: 60–67.

Garthwaite J, Boulton CL (1995) Nitric Oxide Signaling in the Central Nervous System. Annu Rev Physiol 57: 683–706.

Gonzalez-Hernandez T, Rodriguez M (2000) Compartmental organization and chemical profile of dopaminergic and GABAergic neurons in the substantia nigra of the rat. J Comp Neurol 421: 107–135.

Gotti S, Sica M, Viglietti-Panzica C, Panzica G (2005) Distribution of nitric oxide synthase immunoreactivity in the mouse brain. Microsc Res Tech 68: 13–35.

Hnasko TS, Hjelmstad GO, Fields HL, Edwards RH (2012) Ventral tegmental area glutamate neurons: electrophysiological properties and projections. J Neurosci 32: 15076–15085.

Hokfelt T, Rehfeld JF, Skirboll L, Ivemark B, Goldstein M, Markey K (1980) Evidence for coexistence of dopamine and CCK in meso-limbic neurones. Nature 285: 476–478.

Huang PL, Dawson TM, Bredt DS, Snyder SH, Fishman MC (1993) Targeted disruption of the neuronal nitric oxide synthase gene. Cell 75: 1273–1286.

Ibanez-Sandoval O, Tecuapetla F, Unal B, Shah F, Koos T, Tepper JM (2010) Electrophysiological and morphological characteristics and synaptic connectivity of tyrosine hydroxylase-expressing neurons in adult mouse striatum. J Neurosci 30: 6999–7016.

Isaacs KR, Jacobowitz DM (1994) Mapping of the colocalization of calretinin and tyrosine hydroxylase in the rat substantia nigra and ventral tegmental area. Exp Brain Res 99: 34–42.

Klausberger T, Somogyi P (2008) Neuronal diversity and temporal dynamics: The unity of hippocampal circuit operations. Science 321: 53–57.

Klejbor I, Domaradzka-Pytel B, Ludkiewicz B, Wojcik S, Morys J (2004) The relationships between neurons containing dopamine and nitric oxide synthase in the ventral tegmental area. Folia Histochem Cytobiol 42: 83–87.

Klink R, de Kerchove d’Exaerde A, Zoli M, Changeux JP (2001) Molecular and physiological diversity of nicotinic acetylcholine receptors in the midbrain dopaminergic nuclei. J Neurosci 21: 1452–1463.

Knowles RG, Palacios M, Palmer RM, Moncada S (1989) Formation of nitric oxide from L-arginine in the central nervous system: a transduction mechanism for stimulation of the soluble guanylate cyclase. Proc Natl Acad Sci U S A 86: 5159–5162.

Lein ES et al (2007) Genome-wide atlas of gene expression in the adult mouse brain. Nature 445: 168–176.

Liang CL, Sinton CM, German DC (1996) Midbrain dopaminergic neurons in the mouse: co-localization with Calbindin-D28k and calretinin. Neuroscience 75: 523–533.

Mao L, Lau YS, Petroske E, Wang JQ (2001) Profound astrogenesis in the striatum of adult mice following nigrostriatal dopaminergic lesion by repeated MPTP administration. Brain Res Dev Brain Res 131: 57–65.

Matsuda W, Furuta T, Nakamura KC, Hioki H, Fujiyama F, Arai R, Kaneko T (2009) Single nigrostriatal dopaminergic neurons form widely spread and highly dense axonal arborizations in the neostriatum. J Neurosci 29: 444–453.

Mazloom M, Smith Y (2006) Synaptic microcircuitry of tyrosine hydroxylase-containing neurons and terminals in the striatum of 1-methyl-4-phenyl-1,2,3,6-tetrahydropyridine-treated monkeys. J Comp Neurol 495: 453–469.

Meredith GE, Farrell T, Kellaghan P, Tan Y, Zahm DS, Totterdell S (1999) Immunocytochemical characterization of catecholaminergic neurons in the rat striatum following dopamine-depleting lesions. Eur J Neurosci 11: 3585–3596.

Merrill CB, Friend LN, Newton ST, Hopkins ZH, Edwards JG (2015) Ventral tegmental area dopamine and GABA neurons: Physiological properties and expression of mRNA for endocannabinoid biosynthetic elements. Sci Rep 5: 16176.

Mitkovski M, Padovan-Neto FE, Raisman-Vozari R, Ginestet L, da-Silva CA, Del-Bel EA (2012) Investigations into potential extrasynaptic communication between the dopaminergic and nitrergic systems. Front Physiol 3: 372.

Morales M, Margolis EB (2017) Ventral tegmental area: cellular heterogeneity, connectivity and behaviour. Nat Rev Neurosci 18: 73–85.

Morales M, Root DH (2014) Glutamate neurons within the midbrain dopamine regions. Neuroscience 282: 60–68.

Nair-Roberts RG, Chatelain-Badie SD, Benson E, White-Cooper H, Bolam JP, Ungless MA (2008) Stereological estimates of dopaminergic, GABAergic and glutamatergic neurons in the ventral tegmental area, substantia nigra and retrorubral field in the rat. Neuroscience 152: 1024–1031.

O’Byrne MB, Bolam JP, Hanley JJ, Tipton KF (2000) Tyrosine-hydroxylase immunoreactive cells in the rat striatum following treatment with MPP+. Adv Exp Med Biol 483: 369–374.

Olson VG, Nestler EJ (2007) Topographical organization of GABAergic neurons within the ventral tegmental area of the rat. Synapse 61: 87–95.

Omelchenko N, Sesack SR (2009) Ultrastructural analysis of local collaterals of rat ventral tegmental area neurons: GABA phenotype and synapses onto dopamine and GABA cells. Synapse 63: 895–906.

Petroske E, Meredith GE, Callen S, Totterdell S, Lau YS (2001) Mouse model of Parkinsonism: a comparison between subacute MPTP and chronic MPTP/probenecid treatment. Neuroscience 106: 589–601.

Qi J, Zhang S, Wang HL, Barker DJ, Miranda-Barrientos J, Morales M (2016) VTA glutamatergic inputs to nucleus accumbens drive aversion by acting on GABAergic interneurons. Nat Neurosci 19: 725–733.

Rodrigo J, Springall DR, Uttenthal O, Bentura ML, Abadia-Molina F, Riveros-Moreno V, Martinez-Murillo R, Polak JM, Moncada S (1994) Localization of nitric oxide synthase in the adult rat brain. Philosophical Transactions of the Royal Society of London. Series B: Biological Sciences 345: 175–221.

Rogers JH (1992) Immunohistochemical markers in rat brain: colocalization of calretinin and calbindin-D28k with tyrosine hydroxylase. Brain Res 587: 203–210.

Root DH, Mejias-Aponte CA, Qi J, Morales M (2014a) Role of glutamatergic projections from ventral tegmental area to lateral habenula in aversive conditioning. J Neurosci 34: 13906–13910.

Root DH, Mejias-Aponte CA, Zhang S, Wang HL, Hoffman AF, Lupica CR, Morales M (2014b) Single rodent mesohabenular axons release glutamate and GABA. Nat Neurosci 17: 1543–1551.

Root DH, Wang HL, Liu B, Barker DJ, Mod L, Szocsics P, Silva AC, Magloczky Z, Morales M (2016) Glutamate neurons are intermixed with midbrain dopamine neurons in nonhuman primates and humans. Sci Rep 6: 30615.

San Sebastián W, Guillén J, Manrique M, Belzunegui S, Ciordia E, Izal-Azcárate A, Garrido-Gil P, Vázquez-Claverie M, Luquin MR (2007) Modification of the number and phenotype of striatal dopaminergic cells by carotid body graft. Brain 130: 1306–1316.

Sanz E, Yang L, Su T, Morris DR, McKnight GS, Amieux PS (2009) Cell-type-specific isolation of ribosome-associated mRNA from complex tissues. Proc Natl Acad Sci U S A 106: 13939–13944.

Seroogy K, Ceccatelli S, Schalling M, Hökfelt T, Frey P, Walsh J, Dockray G, Brown J, Buchan A, Goldstein M (1988) A subpopulation of dopaminergic neurons in rat ventral mesencephalon contains both neurotensin and cholecystokinin. Brain Res 455: 88–98.

Seroogy K, Schalling M, Brené S, Dagerlind Å, Chai SY, Hökfelt T, Persson H, Brownstein M, Huan R, Dixon J, Filer D, Schlessinger D, Goldstein M (1989) Cholecystokinin and tyrosine hydroxylase messenger RNAs in neurons of rat mesencephalon: peptide/monoamine coexistence studies using in situ hybridization combined with immunocytochemistry. Exp Brain Res 74: 149–162.

Stuber GD, Hnasko TS, Britt JP, Edwards RH, Bonci A (2010) Dopaminergic terminals in the nucleus accumbens but not the dorsal striatum corelease glutamate. J Neurosci 30: 8229–8233.

Tan KR, Yvon C, Turiault M, Mirzabekov JJ, Doehner J, Labouebe G, Deisseroth K, Tye KM, Luscher C (2012) GABA neurons of the VTA drive conditioned place aversion. Neuron 73: 1173–1183.

Tandé D, Höglinger G, Debeir T, Freundlieb N, Hirsch EC, François C (2006) New striatal dopamine neurons in MPTP-treated macaques result from a phenotypic shift and not neurogenesis. Brain 129: 1194–1200.

Tashiro Y, Sugimoto T, Hattori T, Uemura Y, Nagatsu I, Kikuchi H, Mizuno N (1989) Tyrosine hydroxylase-like immunoreactive neurons in the striatum of the rat. Neurosci Lett 97: 6–10.

Taylor SR, Badurek S, Dileone RJ, Nashmi R, Minichiello L, Picciotto MR (2014) GABAergic and glutamatergic efferents of the mouse ventral tegmental area. J Comp Neurol 522: 3308–3334.

Tecuapetla F, Patel JC, Xenias H, English D, Tadros I, Shah F, Berlin J, Deisseroth K, Rice ME, Tepper JM, Koos T (2010) Glutamatergic signaling by mesolimbic dopamine neurons in the nucleus accumbens. J Neurosci 30: 7105–7110.

Tepper JM, Tecuapetla F, Koos T, Ibanez-Sandoval O (2010a) Heterogeneity and diversity of striatal GABAergic interneurons. Front Neuroanat 4: 150.

Tepper JM, Tecuapetla F, Koós T, Ibáñez-Sandoval O (2010b) Heterogeneity and diversity of striatal GABAergic interneurons. Front Neuroanat 4.

Unal B, Ibanez-Sandoval O, Shah F, Abercrombie ED, Tepper JM (2011) Distribution of tyrosine hydroxylase-expressing interneurons with respect to anatomical organization of the neostriatum. Front Syst Neurosci 5: 41.

Unal B, Shah F, Kothari J, Tepper JM (2015) Anatomical and electrophysiological changes in striatal TH interneurons after loss of the nigrostriatal dopaminergic pathway. Brain Struct Funct 220: 331–349.

van Zessen R, Phillips JL, Budygin EA, Stuber GD (2012) Activation of VTA GABA neurons disrupts reward consumption. Neuron 73: 1184–1194.

Vincent SR, Kimura H (1992) Histochemical mapping of nitric oxide synthase in the rat brain. Neuroscience 46: 755–784.

Vong L, Ye C, Yang Z, Choi B, Chua S, Jr., Lowell BB (2011) Leptin action on GABAergic neurons prevents obesity and reduces inhibitory tone to POMC neurons. Neuron 71: 142–154.

Wang HL, Qi J, Zhang S, Wang H, Morales M (2015) Rewarding effects of optical stimulation of ventral tegmental area glutamatergic neurons. J Neurosci 35: 15948–15954.

Wang Y, Marsden PA (1995) Nitric oxide synthases: gene structure and regulation. Adv Pharmacol 34: 71–90.

Xenias HS, Ibanez-Sandoval O, Koos T, Tepper JM (2015) Are striatal tyrosine hydroxylase interneurons dopaminergic? J Neurosci 35: 6584–6599.

Yamaguchi T, Qi J, Wang HL, Zhang S, Morales M (2015) Glutamatergic and dopaminergic neurons in the mouse ventral tegmental area. Eur J Neurosci 41: 760–772.

Yamaguchi T, Sheen W, Morales M (2007) Glutamatergic neurons are present in the rat ventral tegmental area. Eur J Neurosci 25: 106–118.

Yamaguchi T, Wang HL, Li X, Ng TH, Morales M (2011) Mesocorticolimbic glutamatergic pathway. J Neurosci 31: 8476–8490.

Yamaguchi T, Wang HL, Morales M (2013) Glutamate neurons in the substantia nigra compacta and retrorubral field. Eur J Neurosci 38: 3602–3610.

Yoo JH, Zell V, Gutierrez-Reed N, Wu J, Ressler R, Shenasa MA, Johnson AB, Fife KH, Faget L, Hnasko TS (2016) Ventral tegmental area glutamate neurons co-release GABA and promote positive reinforcement. Nat Commun 7: 13697.

Zhang S, Qi J, Li X, Wang HL, Britt JP, Hoffman AF, Bonci A, Lupica CR, Morales M (2015) Dopaminergic and glutamatergic microdomains in a subset of rodent mesoaccumbens axons. Nat Neurosci 18: 386–392.

